# Normal development and fertility of Fut1, Fut2, and Sec1 triple knockout mice

**DOI:** 10.1101/615070

**Authors:** Jiaxi Chen, Zhipeng Su, Chunlei Zhang, Fenge Li, Patrick Hwu, Zhen Wang, Yanping Wang, Yunsen Li, Jiao Tong, Chunchao Chen, Dapeng Zhou

## Abstract

The fucose alpha(1,2) galactose structure (H antigen) is synthesized by α1,2 fucosyltransferases Fut1, Fut2 and Sec1. H antigen has been reported to be involved in cancer progression, neurite migration, synaptic plasticity, host-microbe interaction, blastocyst implantation, and the maintenance of gut microbiome. Genetic depletion of Fut1 or Fut2 only cause defects of alpha1,2 fucosylation in limited tissues because of enzyme redundancy. In this study, we generated mice with deficiencies in Fut1, Fut2, and Sec1 genes to deplete H antigen through BAC Engineering for the generation of ES Cell-Targeting construct. The homogenous triple knockout mice showed no significant decrease of viability or development. Mass spectrometry and Western blot analysis confirmed the absence of H blood group antigen in multiple organs. These results indicate normal development and fertility of mice devoid of blood group H. The fine pathophysiological alterations in these mice remain to be carefully studied, and they may serve as valuable tools to study gut microbiome and host-microbe interactions.

## Introduction

The function of human blood group ABO is a mystery during evolution. Autoantibodies to blood group A, B and H are absent in neonates and reach their maximum at the age of 5 to 10 years (1–2). It is hypothesized that these natural antibodies are induced by gut microbial carbohydrates including lipopolysaccharide structures, while their function remain to be studied. The biosynthesis of blood group ABO structures is intiated by α-2-fucosyltransfearse (EC: 2.4.1.3441). Three members have been cloned for mouse α-2-fucosyltransfearse gene family. Enzyme activities were reported for mouse Fut1 and Fut2, while no enzyme activity has been reported for Sec1(3–4).

α1,2 fucosylated glycans have been reported to be involved in multiple physiological and pathological processes (5). Lai et al. reported that Fut1 and Fut2 were upregulated in breast cancer (6–7). High expression of Fut1 was also reported in small cell lung cancer (8), leukemia (9), ovarian cancer (10–11), prostate cancer (12–13) and hepatocellular carcinoma (14). Mass spectrometry analysis revealed the higher level of expression of α1,2 fucosylated structures including Globo-H in primary hepatocellular carcinoma (15). However, in melanoma samples, Fut1 was reported to be down-regulated (16). Also, contradicting results have been reported on the role of Fut1 in tumor angiogenesis (17–18).

Commensal bacteria induce the expression of α1,2 glycans in epithelial cells (19). Epithelial fucose is used as a dietary carbohydrate by many bacteria. The exact function of α1,2 glycans of gut epithelia is being explored. Defect of intestinal fucosylation in Fut2^−/−^ mice led to increased susceptibility to infection by Salmonella typhimurium. It has also been reported that non-sector status is related to increased susceptibility to Crohn’s disease (20–22).

Both Fut1 and Fut2 have been reported to be expressed in mouse brain and regulate neurite migration and synaptic plasticity. The underlying mechanism appears to be extremely complexed as multiple glycoproteins are modified by α1,2 glycans. Genetic ablation of either Fut1 or Fut2 in mice did not cause complete defect of α1,2 glycans due to redundant expression in mouse brain (23).

The Fut1 and Fut2 gene are located in the same chromosome with a 30 kb distance (Figure 1). This short genetic distance prevents the homologous recombination process required for generating double knockout mice. In this study, we sequentially depleted Fut1 gene and Fut2/Sec1 gene in ES cells and generated triple knockout mice with total deficiency of α1,2-fucosyltransferase enzyme.

**Figure 1:**
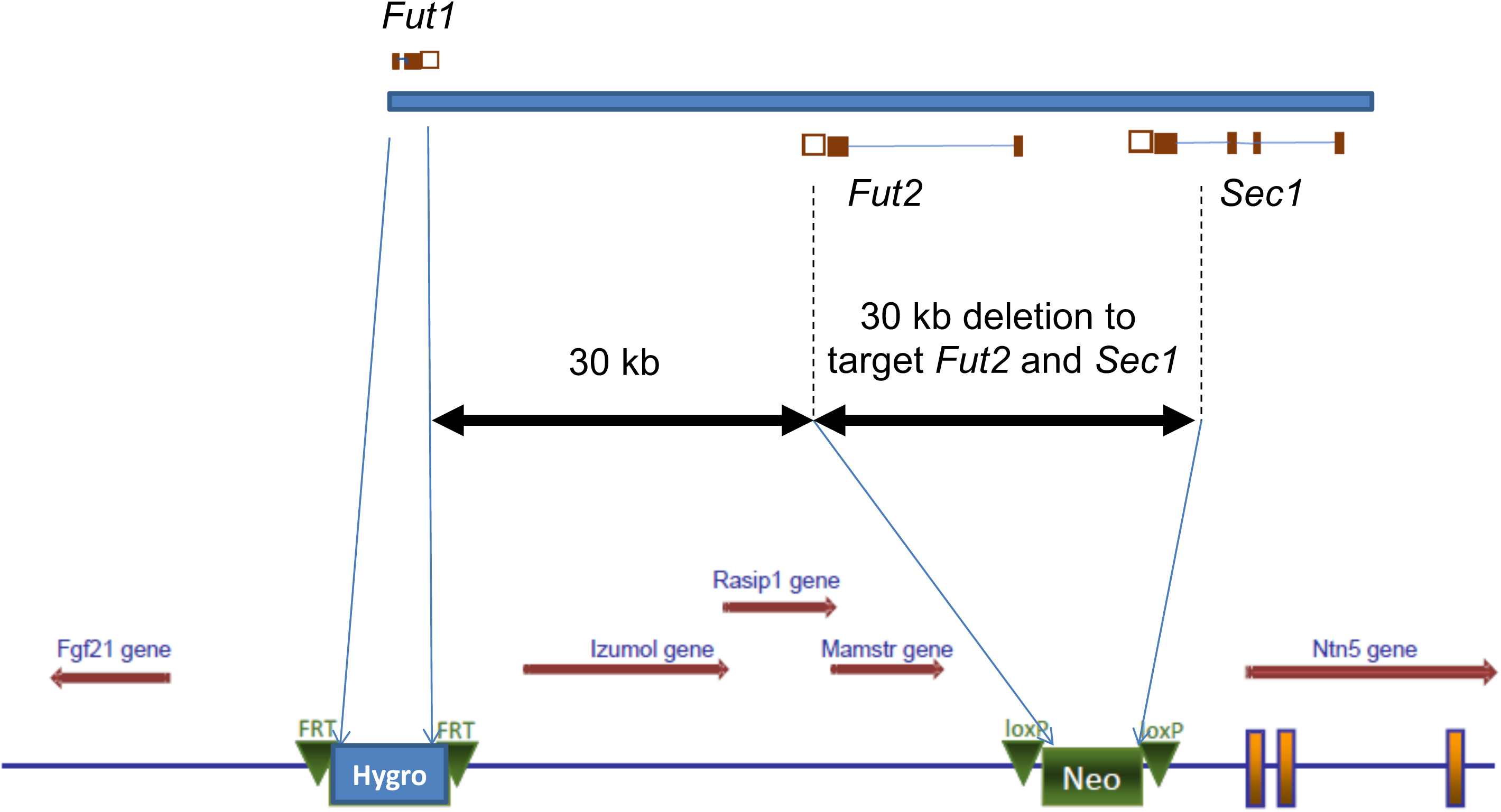
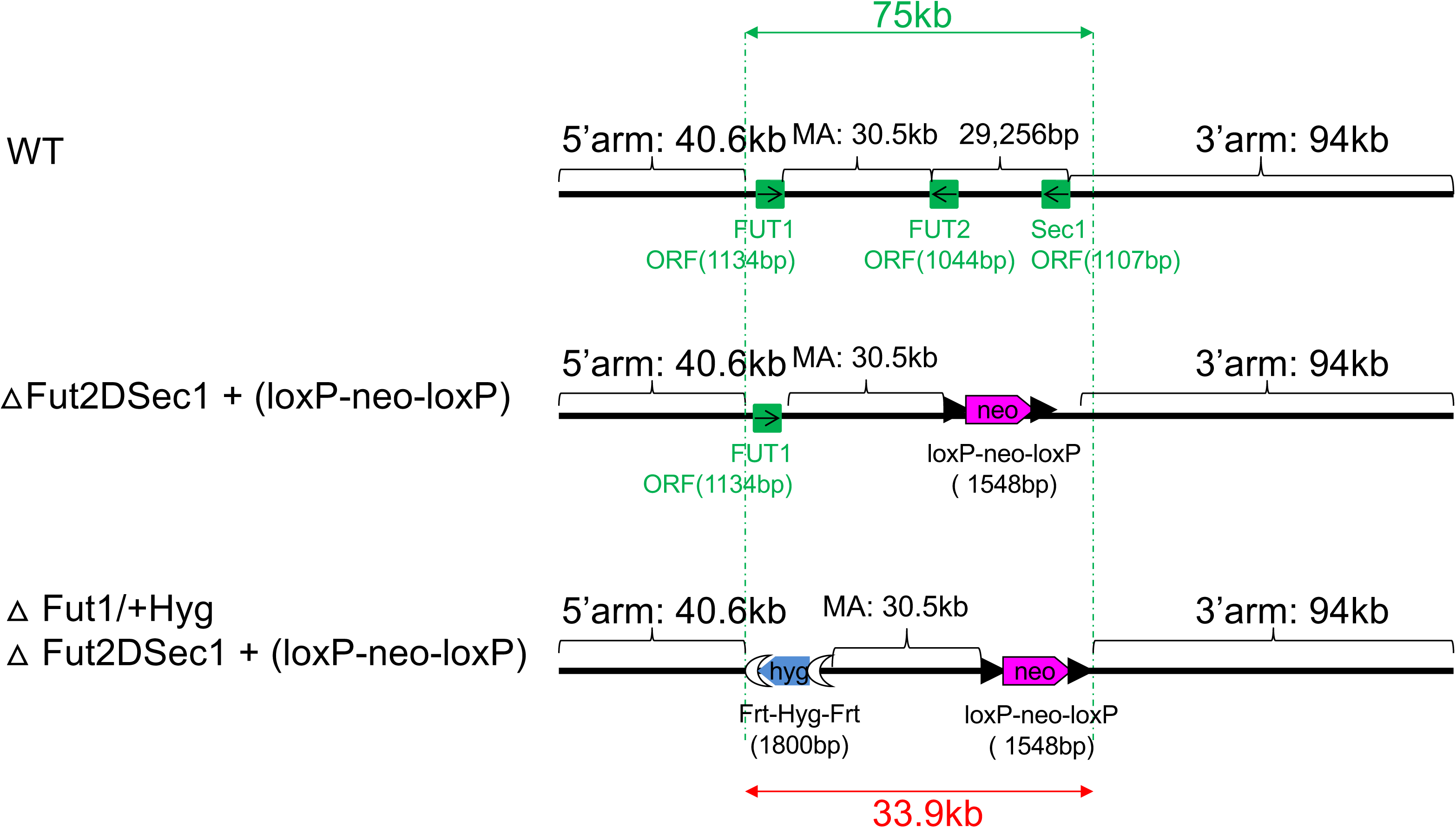
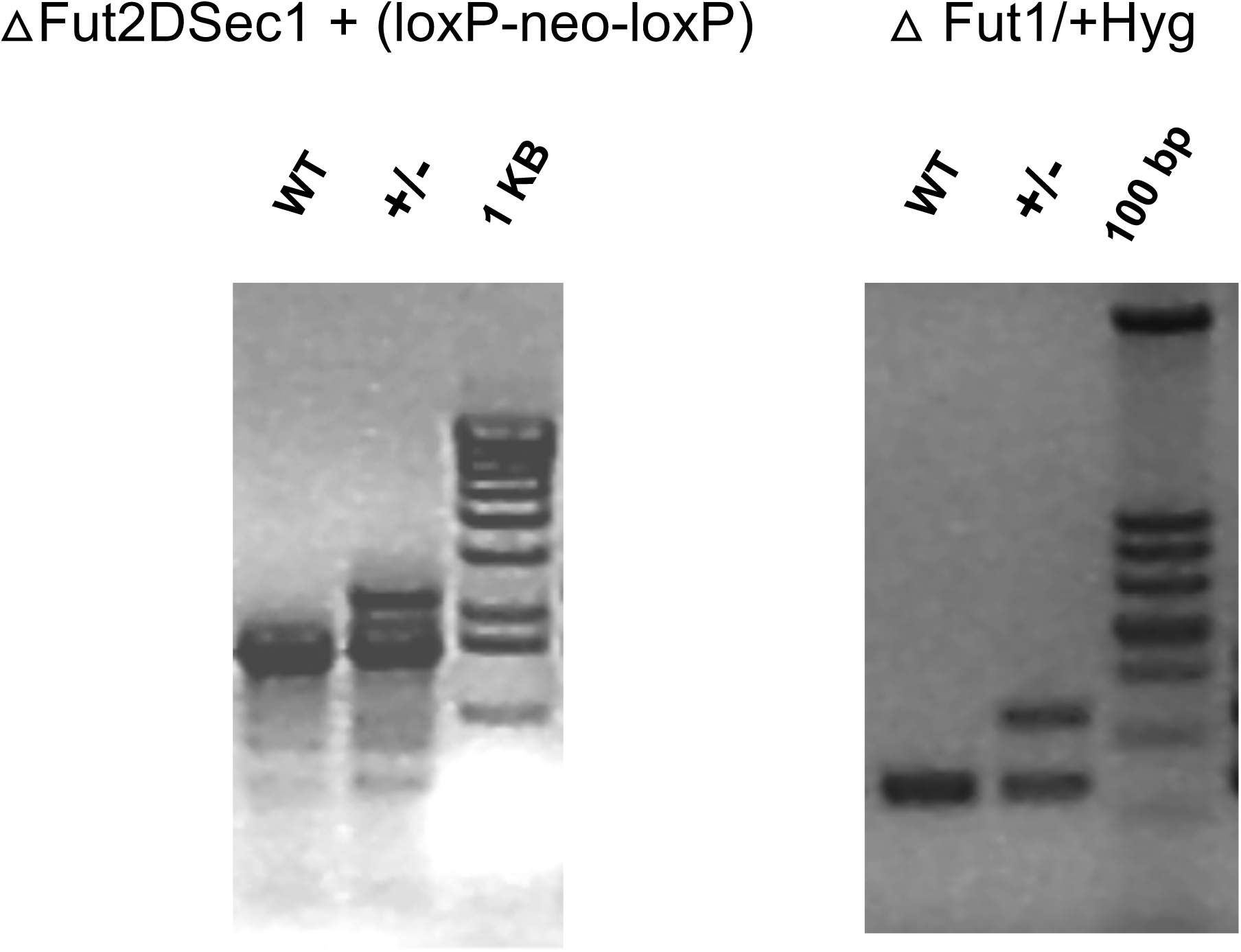
Generation of Fut1/Fut2/Sec1 triple KO mice. A. Two step strategy to replace the protein coding regions (marked as empty boxes) of Fut1, Fut2, and Sec1 by targeting constructs containing neomycin and hygromycin selection markers; B. Scheme of genetic locus for wild type ES cell line, ΔFut2DSec1 + (loxP-neo-loxP) ES cell line, and Δ Fut1/+Hyg Δ Fut2DSec1 + (loxP-neo-loxP) ES cell line. C. PCR results of the product lengths of wildtype allele, Δ Fut1/+Hyg allele, and ΔFut2DSec1 + (loxP-neo-loxP) allele.

## Methods

### Engineering of Bacmid

The Fut1, Fut2, and Sec1 loci are located in chromosome 7 in a 77 kb region (Table 1). A two-step knockout strategy was used to generate the mice with protein-coding regions of all three genes mutated. Firstly, a LoxP-neomycin-LoxP cassette was used to replace a 30 kb region which contains protein coding regions of both Fut2 and Sec1. Secondly, a FRT-hygromycin-FRT cassette was used to replace the protein coding region of Fut1. BAC clone RP23-223G1 was purchased from Children’s Hospital Oakland Research Institute BACPAC Resources Center, and mutated constructs were made by Red/ET recombination technology (24).

**Table 1.** Location of Fut1, Fut2 and Sec1 in mouse genome.

| Gene | Chromosome | Location | Centimorgan | Strand |
| --- | --- | --- | --- | --- |
| Fut1 | 7 | 45617289- 45621059 | 29.39 | + |
| Fut2 | 7 | 45648591- 45666394 | 29.41 | - |
| Sec1 | 7 | 45677686-45694402 | 29.43 | - |

### Generation of Fut1/Fut2/Sec1 triple KO mice

Mutated constructs were electroporated into iTL BA1 (C57BL/6 × 129/SvEv) by Ingenious Targeting Laboratory, NY. Hybrid embryonic stem cells and ES clones bearing mutated loci were selected by neomycin and hygromycin resistance sequentially. Targeted iTL BA1 (C57BL/6 × 129/SvEv) hybrid embryonic stem cells were microinjected into C57BL/6 blastocysts. Resulting chimeras with a high percentage agouti coat color were mated to wild-type C57BL/6N mice to generate F1 heterozygous offspring. Tail DNA was analyzed as described below from pups with agouti or black coat color. Primers F1385hygS and R1385hygS (Table 2, Supplemental Figure 1) were used to determine the mutation of Fut1 locus (replaced by FRT-hygromycin-FRT cassette). Primers Wt1, R1385neoS and N2 were used to determine the mutation of Fut2 and Sec1 locus (replaced by LoxP-neomycin-LoxP cassette).

**Table 2.**
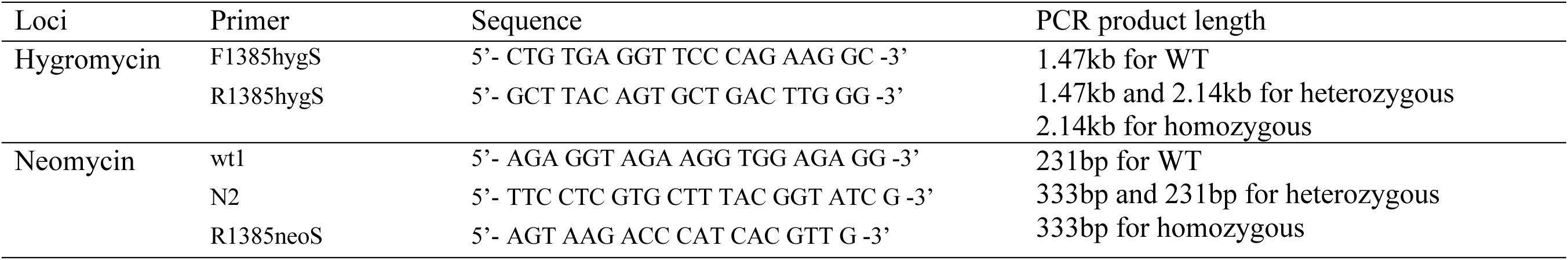
Primers used for genotyping.

### ELISA measurement of natural antibodies to blood group H, A, and B

The biotinylated glycans were provided by Drs. Ola Blixt and James Paulson at the Scripps Research Institute. Biotinylated glycans (1 μg/ml) were bound to streptavidin-coated plates (2 μg/ml), and incubated with serially diluted serum for 2 hours. Binding of glycan-specific IgM and IgG was visualized by a secondary antibody (goat anti-mouse IgM, or goat anti-mouse IgG, Southern Biotech, Birmingham, AL) followed by colorimetric detection. One percent bovine serum albumin was used as blank for determining the cutoff value.

### Generation of a monoclonal antibody specific for blood group H

Fut1/Fut2/ Sec1 triple KO mice were immunized by MCF-7 cell line which express high abundance of blood group H. Monoclonal antibodies were generated by screening against Fuc α1,2 Gal-BSA using ELISA. Fuc α1,2 Gal disaccharide conjugated to BSA (V Labs, UK) was coated to 96 well Costar plates (Corning, NY) at 50 μl per well with a concentration of 5 μg/ml. The plates were blocked by 1% BSA (Sigma, St. Louis), and the antibody containing sera were added after being serially diluted. After 1 hour of incubation, the plates were washed 5 times with PBS/0.05% Tween, and HRP-conjugated donkey anti-mouse IgG antibody (Jackson Immuno Reseach, TN) was added. The bound secondary antibody was detected by a HRP substrate kit from Pharmingen.

In Fut1/Fut2/Sec1 triple KO mice vaccinated by MCF cells, serum IgG reactivity to Fuc α1,2 Gal-BSA reached the titer of 1:5,000. 2 of the immunized mice were sacrificed, and the spleenocytes were fused to Sp2/0-Ag14 myeloma cell line, according to a protocol established at the Monoclonal antibody core facility at the University of Texas MD Anderson Cancer Center. The supernatant of hybridomas were screen by ELISA. One clone, 4B, that produce high titer of anti-monoclonal antibodies against Fuc α1,2 Gal-BSA were further sub-cloned twice. This antibody clone was submitted to Consortium of Functional Glycomics glycan-array core facility at Emory University to test its specificity toward 600 mammalian glycans, as well as a glycan-chip for blood group related structures.

### Glycan array analysis for monoclonal antibody 4B

4B antibody was analyzed on a glycan array at Emory Comprehensive Glycomics Core, using standard procedures in Tris Saline buffer at pH 7.4 containing 2 mM CaCl_2_ and 2 mM MgCl_2_ with 1 mg/ml BSA and .05% Tween 20. Antibody at indicated concentration was incubated for 1 hour at room temperature under a coverslip on the CFG array on a microscope slide, washed and the secondary antibody (Alexa647 labeled goat anti-mouse IgG) was similarly applied. The slide was washed with buffer, dried, and scanned at 647nm.

### Western blot analysis

Protein concentration was determined by the Bradford method and proteins were separated by SDS-PAGE gel electrophoresis. Proteins were transferred onto PVDF membranes (Millipore, Burlington, MA, USA). The blots were blocked for 2 h at room temperature with 5% non-fat milk in PBS/0.05% Tween 20 and then incubated with the primary antibody (1 µg/mL) overnight at 4 °C. After washing, the PVDF membrane was incubated with horseradish peroxidase (HRP)-conjugated secondary antibody at room temperature for 1 h. The transferred proteins were visualized with chemiluminescence detection kit (Millipore).

### Mass spectrometry analysis

Glycosphingolipids were extracted and permethylated as described (25–26). Qualitative analysis of fucosylated neutral GSLs was performed in the positive ion mode on the LTQ-XL mass spectrometer (Thermo Fischer Scientific, Waltham, MA, USA) by using a metal needle for direct infusion of samples dissolved in methanol, with a flow rate of 5 μl/min and at ion spray voltage 3.5 kV, capillary voltage 35 V, capillary temperature 350°C, injection time 100 ms, activation time 30 ms, and isolation width m/z 1.5. All ions were detected as sodium adducts. To gather as much structure information as possible, MS^3^, MS^4^, and MS^5^ spectra of all the fucosylated neutral GSLs were obtained as well.

### Metagenomic analysis of microbiome

Fecal DNA from mouse colon was extracted by QIAamp PowerFecal DNA Kit (Qiagen, Germany) with minor modifications. Samples were processed using Minibeadbeater-16 (Biospec, USA) at maximum speed for 1min instead of vortex, and other steps were according to the user manual. DNA concentration the was measured by a NanoDrop 2000 (Thermo Scientific) and stored at −20C°. All DNA samples were fragmented by Bioruptor sonicator and the libraries were prepared using NEXTflex Rapid DNA-Seq Kit (Bioo Scientific, Austin, Texas, USA) with an average insert size of 450 bp. All liberaries were sequenced on Illumina HiSeq X Ten platform using the 150 bp paired-end module.

Raw reads were trimmed, filtered and aligned against a custom database to screen for human contamination. MetaPhlAn (v2.0) (27) was applied to determine the relative abundance of bacterial species present in all samples. SOAPdenovo (28) and MetaGeneMark (29) software were used to perform de novo assembly and gene prediction with the high quality reads, respectively. All predicted genes were aligned by CD-HIT (identity >95% and coverage >90%) (30) to get the non-redundant gene catalogue. To get the relative gene abundance for each gene, the high quality reads from each sample were aligned against the gene catalogue by SOAP2 (identity > 95%). We aligned putative amino acid sequences from the gene catalogue against VFDB, CAZy, eggNOG and KEGG databases (release 59.0) using BLASTP (e-value ⩽1e-5). The Metastats method was used to evaluate abundance features between groups (http://metastats.cbcb.umd.edu/detection.html). Linear discriminant analysis (LDA) effect size (LEfSe) method was also used to identify species that show statistically significant differential abundances among groups (31). Heat maps and hierarchical clustering was generated by using R package “heatmap” and “stats” respectively. The effect size R and statistical significance p were determined by ANalysis Of SIMilarity (ANOSIM), assessing statistical significance with 9999 permutations.

## Results and Discussion

### Normal viability and development of Fut1/Fut2/Sec1 triple KO mice

Triple knockout mice with deletion of Fut1, Fut2, and Sec1 were genotyped by PCR (Figure 1, Table 2). The homozygous mutant of triple KO mice showed no statistically significant changes in viability as compared to wild type or heterozygous littermate control (Table 3). No significant loss of body weight or other abnormal development was observed.

**Table 3.** Viablity of Fut1/Fut2/Sec triple KO mice.

| Gender | WT(+/+) | Heterozygous(+/-) | KO(-/-) | Total screened |
| --- | --- | --- | --- | --- |
| ♀ | 16.9% | 25.4% | 3.4% | 27 |
| ♂ | 15.3% | 30.5% | 8.5% | 32 |
| Total screened | 32.2% | 55.9% | 11.9% | 59 |

### Generation of a monoclonal antibody specific to type II H and Lewis y glycan structures in Fut1/Fut2/Sec1 triple KO mice

A hybridoma 4B was isolated from MCF-7 immunized mice, and further sub-cloned to generate monoclonal antibody. 600-glycan array analysis showed that the antibody binds with strong affinity to multiple glycans with Fuc α2 Gal β4 GlcNAc β at non-reducing ends (Figure 2A, Supplemental Table 1). 4B also binds to glycans with Lewis y structure, Fuc α2 Gal β4 (Fuc α3) GlcNAc β, at non-reducing ends. We further tested 18 glycans including 6 types of blood group H, A and B structures (Figure 2B, Supplemental Table 2). 4B binds to H2 (Fuc α2 Gal β4 GlcNAc β), H5 (Fuc α2 Gal β3 Ga β) and H6 (Fuc α2 Gal β4 Glc β) structures, but not H1 (Fuc α2 Gal β3 GlcNAc β), H3 (Fuc α2 Gal β3 GalNAc α), or H4 (Fuc α2 Gal β3 GalNAc β) structures. B4 does not bind to Globo-H structure (Fuc α2 Gal β3 GalNAc β Gal α4 Gal β4 Glc β) either by the 600-glycan array.

**Figure 2:**
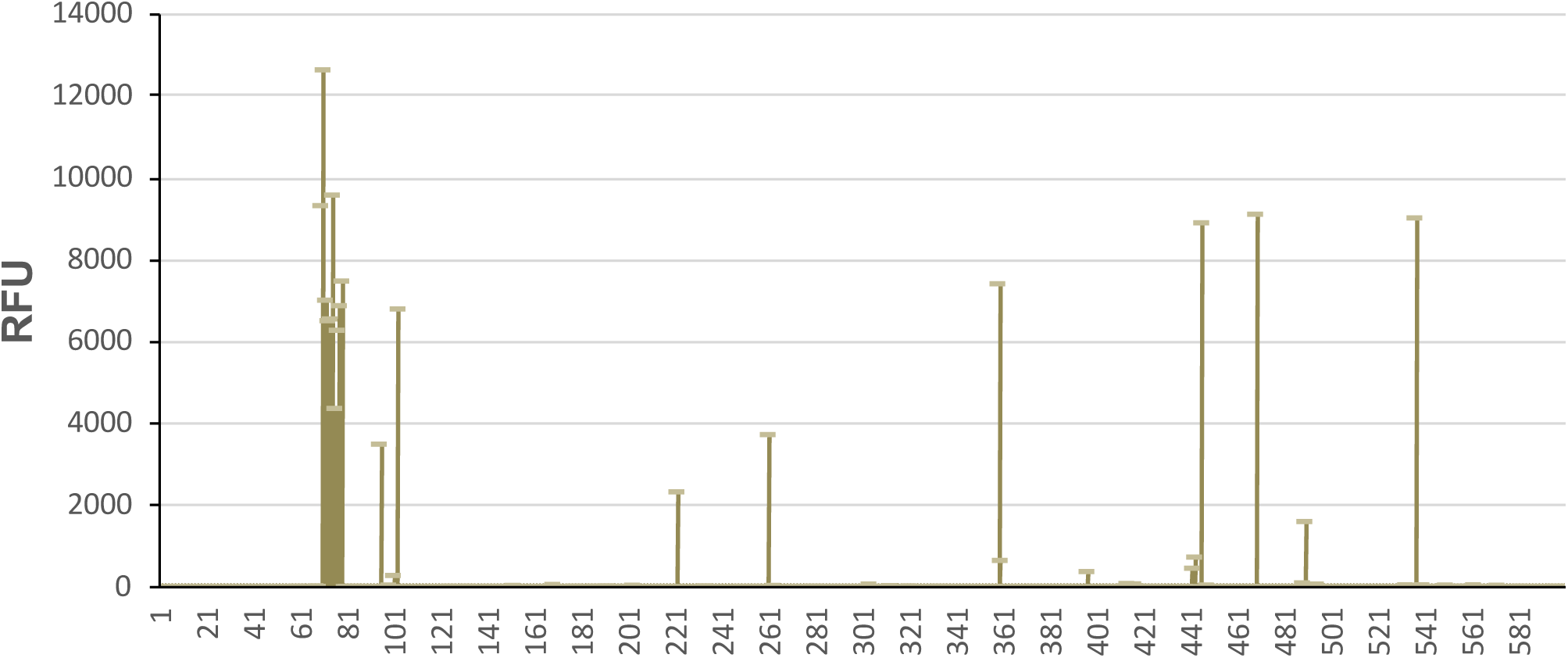
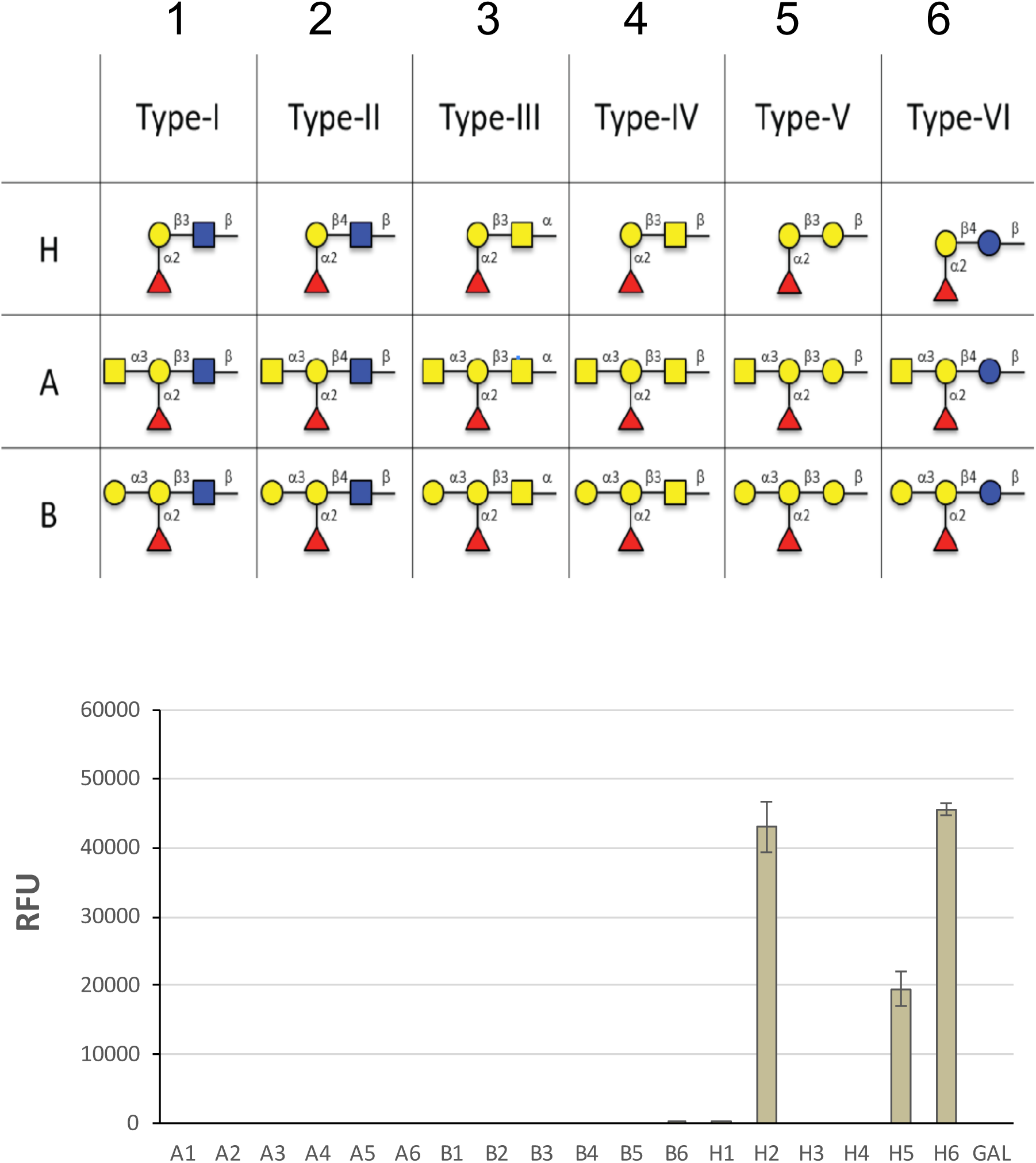
Glycan array analysis of a blood-group-H-specific monoclonal antibody generated in Fut1/Fut2/Sec1 triple KO mice. A. Binding of 4B antibody to 600 glycans; B. Binding of 4B antibody to H type of antigens containing α1,2 fucose at non-reducing end. Data are presented as RFU (relative fluorescence units) vs. Glycan ID. The glycan structures ordered as they appear on the array are shown on the left panel of the spread sheet. The histogram is shown on the center and the glycans ordered according to RFU (highest to lowest) are on the right of the histogram.

### Lack of H blood group antigen in Fut1/Fut2/Sec1 triple KO mice

We tested the presence of H blood group antigen by Western blot analysis of gastric, intestine and colon tissues using 4B antibody. As compared to Fut1 KO mice or Fut2 KO mice, the Fut1/Fut2/Sec1 triple KO mice showed complete lack of H blood group expression (Figure 3A). The lack of Fuc α2 Gal structure was further confirmed by mass spectrometry analysis of neutral glycosphingolipids purified from testis (Figure 3B).

**Figure 3:**
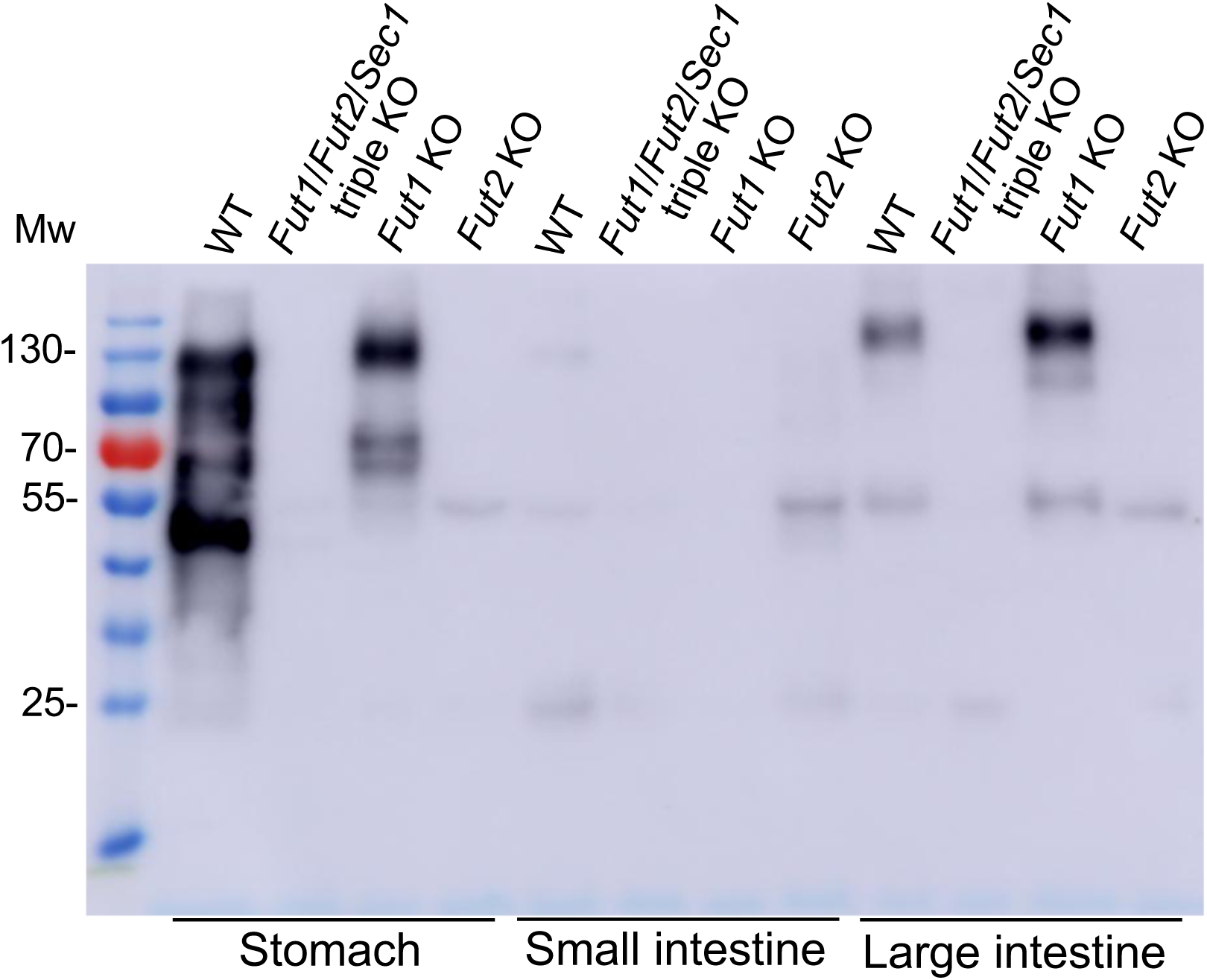
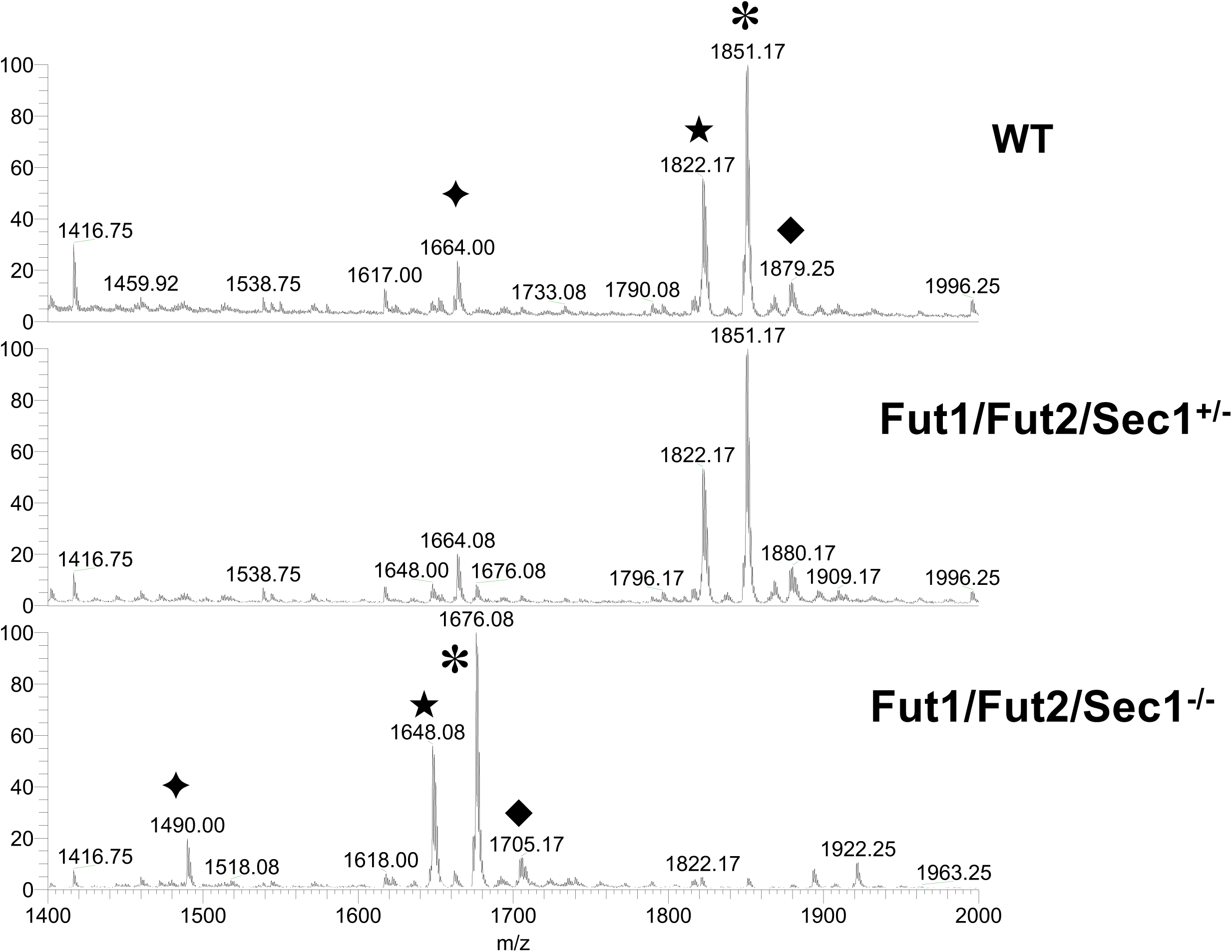
Lack of blood group antigens in Fut1/Fut2/Sec1 triple KO mice. A. Western blot analysis of stomach, small intestine, and large intestine of Fut1/Fut2/Sec1 triple KO, Fut1 KO, and Fut2 KO mice; B. Mass spectrometry analysis of Fut1/Fut2/Sec1^−/−^, Fut1/Fut2/Sec1^+/−^, and WT mice. Symbols indicate the pair of ions with m/z difference of 174, suggesting fucosylated glycosphingolipid structures which are absent in Fut1/Fut2/Sec1^−/−^ mice.

### Normal serum reactivity to blood group ABO in Fut1/Fut2/Sec1 triple KO mice

We measured the titer of natural antibodies to blood group ABO antigens in Fut1/Fut2/Sec1 triple KO mice and wild type mice. No significant differences of either IgM or IgG were found in 2-month old mice in natural anti-ABO antibodies (Figure 4). We further performed metagenomic analysis of microbiome of colon from Fut1/Fut2/Sec1 triple KO or wild type littermate control. No significant difference of bacteria species and abundance were observed either (Supplemental Figure 2).

**Figure 4:**
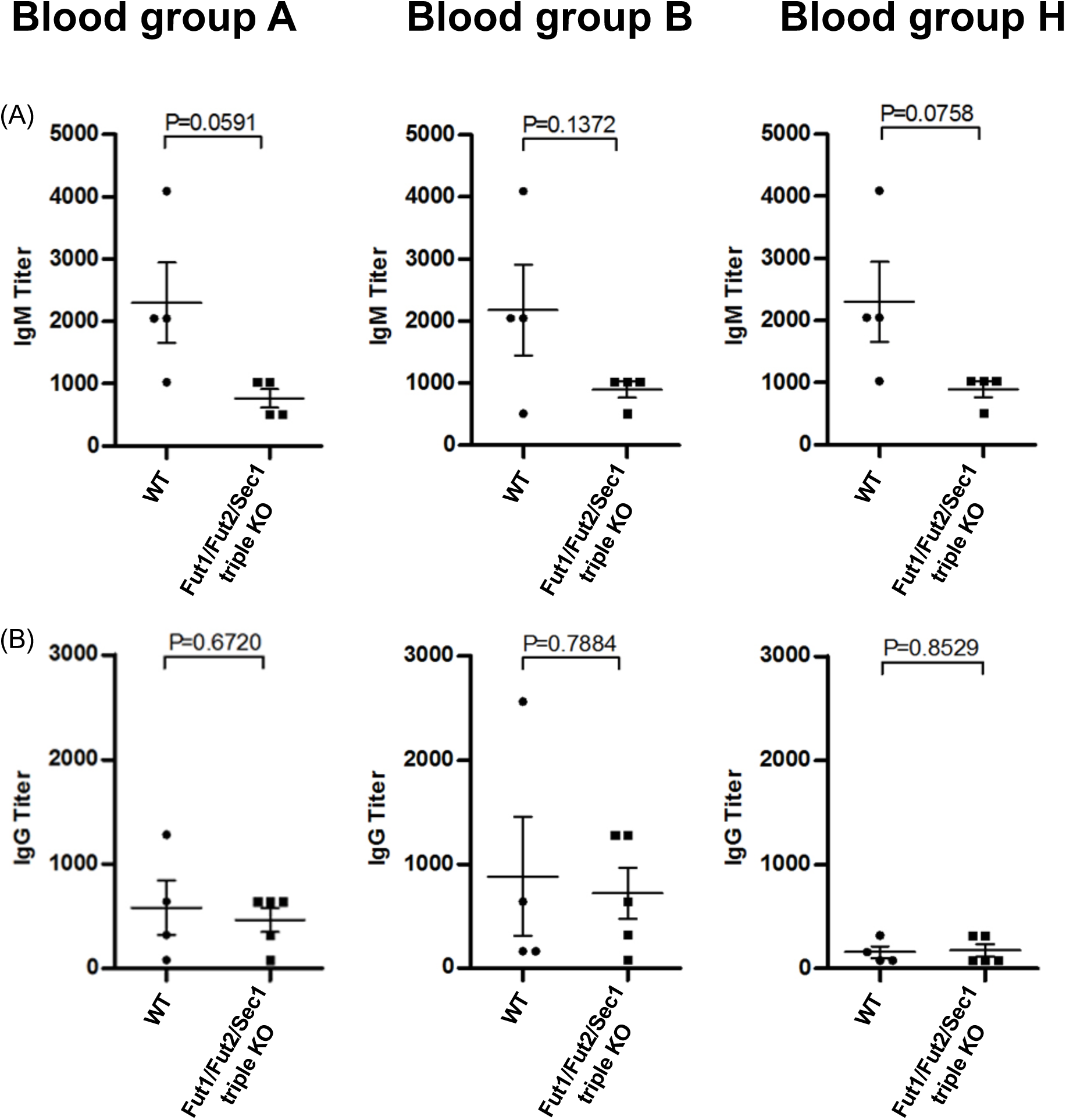
Antibody titers to blood group H, A, and B in in Fut1/Fut2/Sec1 triple KO mice, Fut1 KO, Fut2 KO, and WT mice. Natural antibodies to blood group sugar H, A and B were measured by ELISA as described.

In summary, mice devoid of blood group H showed normal development and fertility. The fine pathophysiological alterations in these mice remain to be carefully studied, such as cancer progression, neurite migration, and synaptic plasticity. The Fut1/Fut2/Sec1 triple KO mice may also be a tool to study the role of endogenous fucosylated glycans in gut-microbiome interaction with complete lack of H blood group expression.

## Abbreviations

Fut1: α-2 fucosyltransferase I
Fut2: α-2 fucosyltransferase II
Sec1: α-2 fucosyltransferase III
Blood group H: Fuc α2 Gal β
Blood group A: GalNAc α3 Fuc α2 Gal β
Blood group B: Gal α3 Fuc α2 Gal β
CFG: consortium of functional glycomics
Lewis y: Fuc α2 Gal β4 (α3) GlcNAc β
Type I H: Fuc α2 Gal β3 GlcNAc β
Type II H: Fuc α2 Gal β4 GlcNAc β
Type III H: Fuc α2 Gal β3 GalNAc α
Type IV H: Fuc α2 Gal β3 GalNAc β
Type V H: Fuc α2 Gal β3 Ga β
Type VI H: Fuc α2 Gal β4 Glc β

## Authors’ Contributions

Dapeng Zhou designed this study. Jiaxi Chen, Chunlei Zhang, Zhipeng Su, Fenge Li, Patrick Hwu, Zhen Wang, Yanping Wang, Yunsen Li, Jiao Tong, Chunchao Chen, Dapeng Zhou contributed to the collection, analysis and interpretation of data. Dapeng Zhou wrote the manuscript. All authors read and approved the final manuscript.

## Acknowledgments

We thank Drs. Xuezheng Song and David Smith at Emory Comprehensive Glycomics Core. This work was supported by National Key Research and Development Plan grant 2017YFA0505901, Fundamental Research Funds for the Central Universities 22120180201, and National Natural Science Foundation of China grant 31870972. All these sponsors have no roles in the study design, or the collection, analysis, and interpretation of data.

## Supplemental online materials

**Supplemental Figure 1:** sites of PCR primers used for genotyping of Δ Fut1/+Hyg allele, and ΔFut2DSec1 + (loxP-neo-loxP) allele.

**Supplemental Figure 2:** metagenomic analysis of microbiome of colon from Fut1/Fut2/Sec1 triple KO or wild type littermate control.

**Supplemental Table 1:** binding of 4B antibody to 600 glycans.

**Supplemental Table 2:** binding of 4B antibody to H type of antigens containing a1,2 fucose at non-reducing end.

## Reference

1. Auf der Maur C, Hodel M, Nydegger UE, Rieben R. Age dependency of ABO histo-blood group antibodies: reexamination of an old dogma. Transfusion. 1993 Nov-Dec;33(11):915–8.

2. Rieben R, Buchs JP, Flückiger E, Nydegger UE. Antibodies to histo-blood group substances A and B: agglutination titers, Ig class, and IgG subclasses in healthy persons of different age categories. Transfusion. 1991 Sep;31(7):607–15.

3. Domino SE, Zhang L, Lowe JB. Molecular cloning, genomic mapping, and expression of two secretor blood group alpha (1,2)fucosyltransferase genes differentially regulated in mouse uterine epithelium and gastrointestinal tract. J Biol Chem. 2001 Jun 29;276(26):23748–56.

4. Lin B, Saito M, Sakakibara Y, Hayashi Y, Yanagisawa M, Iwamori M. Characterization of three members of murine alpha1,2-fucosyltransferases: change in the expression of the Se gene in the intestine of mice after administration of microbes. Arch Biochem Biophys. 2001 Apr 15;388(2):207–15.

5. Schneider M, Al-Shareffi E, Haltiwanger RS. Biological functions of fucose in mammals. Glycobiology. 2017 Jul 1;27(7):601–618.

6. Lai TY, Chen IJ, Lin RJ, Liao GS, Yeo HL, Ho CL, Wu JC, Chang NC, Lee AC, Yu AL. Fucosyltransferase 1 and 2 play pivotal roles in breast cancer cells. Cell Death Discov. 2019 Mar 6;5:74.

7. Chang WW, Lee CH, Lee P, Lin J, Hsu CW, Hung JT, Lin JJ, Yu JC, Shao LE, Yu J, Wong CH, Yu AL. Expression of Globo H and SSEA3 in breast cancer stem cells and the involvement of fucosyl transferases 1 and 2 in Globo H synthesis. Proc Natl Acad Sci U S A. 2008 Aug 19;105(33):11667–72.

8. Tokuda N, Zhang Q, Yoshida S, Kusunoki S, Urano T, Furukawa K, Furukawa K. Genetic mechanisms for the synthesis of fucosyl GM1 in small cell lung cancer cell lines. Glycobiology. 2006 Oct;16(10):916–25.

9. Cao Y, Merling A, Karsten U, Schwartz-Albiez R. The fucosylated histo-blood group antigens H type 2 (blood group O, CD173) and Lewis Y (CD174) are expressed on CD34+ hematopoietic progenitors but absent on mature lymphocytes. Glycobiology. 2001 Aug;11(8):677–83.

10. Liu J, Zheng M, Qi Y, Wang H, Liu M, Liu Q, Lin B. Lewis(y) antigen-mediated positive feedback loop induces and promotes chemotherapeutic resistance in ovarian cancer. Int J Oncol. 2018 Oct;53(4):1774–1786.

11. Li FF, Sha D, Qin XY, Li CZ, Lin B. Alpha1,2-fucosyl transferase gene, the key enzyme of Lewis y synthesis, promotes Taxol resistance of ovarian carcinoma through apoptosis-related proteins. Neoplasma. 2018;65(4):515–522.

12. Nonaka M, Fukuda MN, Gao C, Li Z, Zhang H, Greene MI, Peehl DM, Feizi T, Fukuda M. Determination of carbohydrate structure recognized by prostate-specific F77 monoclonal antibody through expression analysis of glycosyltransferase genes. J Biol Chem. 2014 Jun 6;289(23):16478–86.

13. Fukushima K, Satoh T, Baba S, Yamashita K. alpha1,2-Fucosylated and beta-N-acetylgalactosaminylated prostate-specific antigen as an efficient marker of prostatic cancer. Glycobiology. 2010 Jan;20(4):452–60.

14. Kuo HH, Lin RJ, Hung JT, Hsieh CB, Hung TH, Lo FY, Ho MY, Yeh CT, Huang YL, Yu J, Yu AL. High expression FUT1 and B3GALT5 is an independent predictor of postoperative recurrence and survival in hepatocellular carcinoma. Sci Rep. 2017 Sep 7;7(1):10750.

15. Zhu J, Wang Y, Yu Y, Wang Z, Zhu T, Xu X, Liu H, Hawke D, Zhou D, Li Y. Aberrant fucosylation of glycosphingolipids in human hepatocellular carcinoma tissues. Liver Int. 2014 Jan;34(1):147–60.

16. Agrawal P, Fontanals-Cirera B, Sokolova E, Jacob S, Vaiana CA, Argibay D, Davalos V, McDermott M, Nayak S, Darvishian F, Castillo M, Ueberheide B, Osman I, Fenyö D, Mahal LK, Hernando E. A Systems Biology Approach Identifies FUT8 as a Driver of Melanoma Metastasis. Cancer Cell. 2017 Jun 12;31(6):804–819.e7.

17. Moehler TM, Sauer S, Witzel M, Andrulis M, Garcia-Vallejo JJ, Grobholz R, Willhauck-Fleckenstein M, Greiner A, Goldschmidt H, Schwartz-Albiez R. Involvement of alpha 1-2-fucosyltransferase I (FUT1) and surface-expressed Lewis(y) (CD174) in first endothelial cell-cell contacts during angiogenesis. J Cell Physiol. 2008 Apr;215(1):27–36.

18. Mathieu S, Gerolami R, Luis J, Carmona S, Kol O, Crescence L, Garcia S, Borentain P, El-Battari A. Introducing alpha(1,2)-linked fucose into hepatocarcinoma cells inhibits vasculogenesis and tumor growth. Int J Cancer. 2007 Oct 15;121(8):1680–9.

19. Goto Y, Obata T, Kunisawa J, Sato S, Ivanov II, Lamichhane A, Takeyama N, Kamioka M, Sakamoto M, Matsuki T, Setoyama H, Imaoka A, Uematsu S, Akira S, Domino SE, Kulig P, Becher B, Renauld JC, Sasakawa C, Umesaki Y, Benno Y, Kiyono H. Innate lymphoid cells regulate intestinal epithelial cell glycosylation. Science. 2014 Sep 12;345(6202):1254009.

20. Forni D, Cleynen I, Ferrante M, Cassinotti A, Cagliani R, Ardizzone S, Vermeire S, Fichera M, Lombardini M, Maconi G, de Franchis R, Asselta R, Biasin M, Clerici M, Sironi M. ABO histo-blood group might modulate predisposition to Crohn’s disease and affect disease behavior. J Crohns Colitis. 2014 Jun;8(6):489–94.

21. Parmar AS, Alakulppi N, Paavola-Sakki P, Kurppa K, Halme L, Färkkilä M, Turunen U, Lappalainen M, Kontula K, Kaukinen K, Mäki M, Lindfors K, Partanen J, Sistonen P, Mättö J, Wacklin P, Saavalainen P, Einarsdottir E. Association study of FUT2 (rs601338) with celiac disease and inflammatory bowel disease in the Finnish population. Tissue Antigens. 2012 Dec;80(6):488–93.

22. McGovern DP, Jones MR, Taylor KD, Marciante K, Yan X, Dubinsky M, Ippoliti A, Vasiliauskas E, Berel D, Derkowski C, Dutridge D, Fleshner P, Shih DQ, Melmed G, Mengesha E, King L, Pressman S, Haritunians T, Guo X, Targan SR, Rotter JI; International IBD Genetics Consortium. Fucosyltransferase 2 (FUT2) non-secretor status is associated with Crohn’s disease. Hum Mol Genet. 2010 Sep 1;19(17):3468–76.

23. Wibowo A, Peters EC, Hsieh-Wilson LC. Photoactivatable glycopolymers for the proteome-wide identification of fucose-α(1-2)-galactose binding proteins. J Am Chem Soc. 2014 Jul 9;136(27):9528–31.

24. Bird AW, Erler A, Fu J, Hériché JK, Maresca M, Zhang Y, Hyman AA, Stewart AF. High-efficiency counter selection recombineering for site-directed mutagenesis in bacterial artificial chromosomes. Nat Methods. 2011 Dec 4;9(1):103–9

25. Yin AB, Hawke D, Zhou D. Mass spectrometric analysis of glycosphingolipid antigens. J Vis Exp. 2013 Apr 16;(74).

26. Tahiri F, Li Y, Hawke D, Ganiko L, Almeida I, Levery S, Zhou D. Lack of iGb3 and Isoglobo-Series Glycosphingolipids in Pig Organs Used for Xenotransplantation: Implications for Natural Killer T-Cell Biology. J Carbohydr Chem. 2013 Jan 1;32(1):44–67.

27. Truong DT, Franzosa EA, Tickle TL, Scholz M, Weingart G, Pasolli E, Tett A, Huttenhower C, Segata N: MetaPhlAn2 for enhanced metagenomic taxonomic profiling. Nat Methods 2015, 12:902–903.

28. Luo R, Liu B, Xie Y, Li Z, Huang W, Yuan J, He G, Chen Y, Pan Q, Liu Y, et al: SOAPdenovo2: an empirically improved memory-efficient short-read de novo assembler. Gigascience. 2012 Dec 27;1(1):18. Erratum in: Gigascience. 2015;4:30.

29. Zhu W, Lomsadze A, Borodovsky M: Ab initio gene identification in metagenomic sequences. Nucleic Acids Res. 2010 Jul;38(12):e132.

30. Fu L, Niu B, Zhu Z, Wu S, Li W: CD-HIT: accelerated for clustering the next-generation sequencing data. Bioinformatics 2012, 28:3150–3152.

31. Segata N, Izard J, Waldron L, Gevers D, Miropolsky L, Garrett WS, Huttenhower C: Metagenomic biomarker discovery and explanation. Genome Biol. 2011 Jun 24;12(6):R60.

