## Supplemental Figure 1 for "Normal development and fertility of Fut1, Fut2, and Sec1 triple knockout mice"

### Slide 1
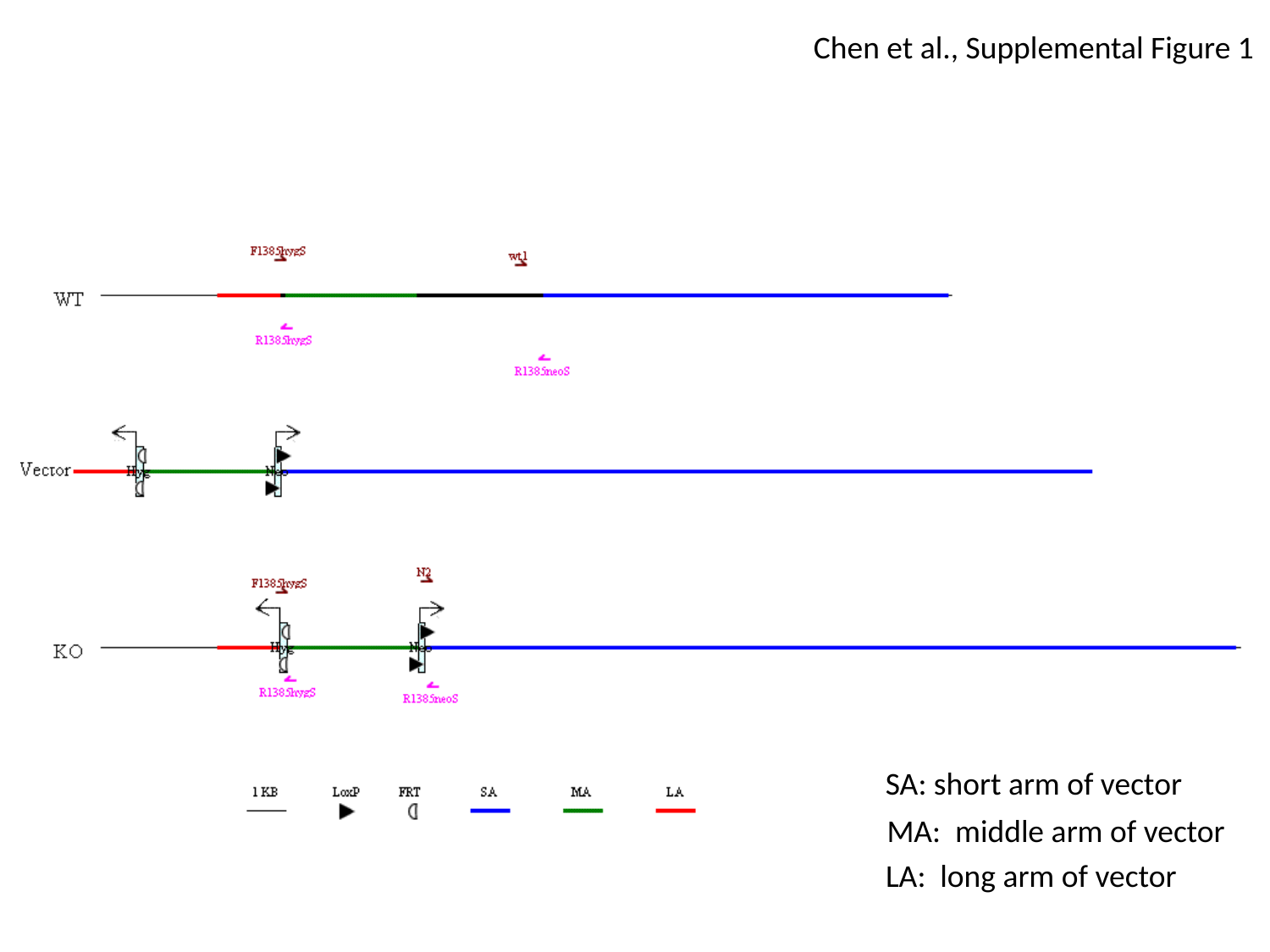

Chen et al., Supplemental Figure 1
SA: short arm of vector
MA: middle arm of vector
LA: long arm of vector
