## Supplementary figures and images for "Normal development and fertility of Fut1, Fut2, and Sec1 triple knockout mice"

### Supplemental Figure 2

#### Slide 1
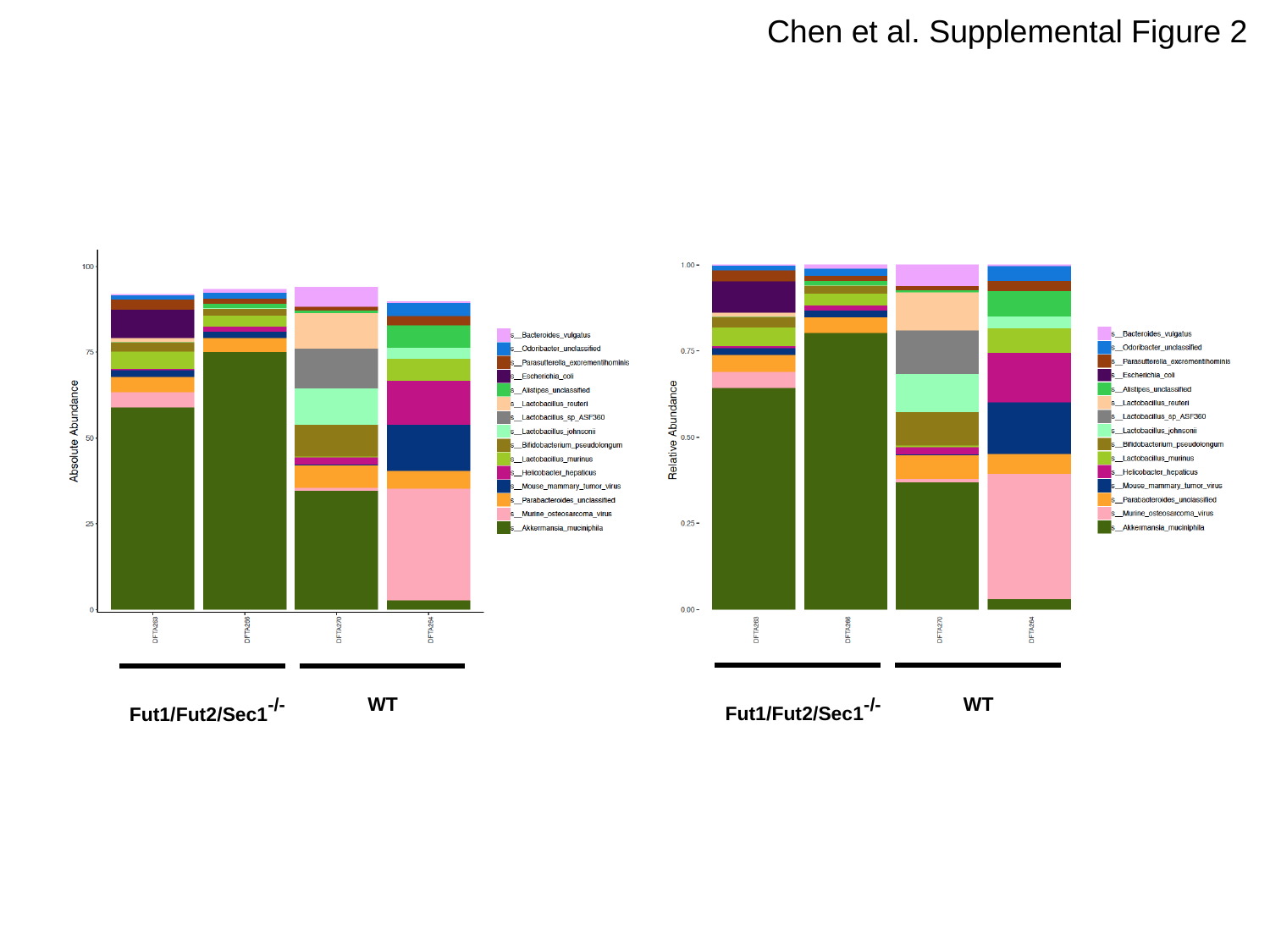

Chen et al. Supplemental Figure 2
Fut1/Fut2/Sec1^-/-^
Fut1/Fut2/Sec1^-/-^
WT
WT
